# Distinct trends in chemical composition of proteins from metagenomes in redox and salinity gradients

**DOI:** 10.1101/2020.04.01.020008

**Authors:** Jeffrey M. Dick, Miao Yu, Jingqiang Tan

## Abstract

Thermodynamic influences on the chemical compositions of proteins have remained enigmatic despite much work that demonstrates the impact of environmental conditions on amino acid frequencies. Here, we show that the dehydrating effect of salinity is apparent in protein sequences inferred from metagenomes and metatranscriptomes. The stoichiometric hydration state (*n*_H_2_O_), derived from the number of water molecules in theoretical reactions to form proteins from a particular set of basis species (glutamine, glutamic acid, cysteine, O_2_, H_2_O), decreases along salinity gradients including the Baltic Sea, Amazon River and ocean plume, and other samples from freshwater and marine environments. Analysis of other metagenomic datasets shows that differences in carbon oxidation state, rather than stoichiometric hydration state, are a stronger indicator of redox gradients than salinity gradients. These compositional metrics can help to identify thermodynamic effects in the distribution of proteins along chemical gradients at scales from geologic systems to cells.

## Introduction

How microbial communities adapt to environmental gradients is a major challenge at the intersection of geochemistry, microbiology, and biochemistry. Patterns of amino acid usage in proteins are important indicators of microbial adaptation, and amino acid composition at the genome level is well known to depend on growth temperature^1^. Furthermore, measures of evolutionary distance and community composition based on protein sequences predicted from metagenomic sequencing are strongly associated with environmental temperature and pH^2^. It is widely held that the effect of amino acid substitutions on the structural stability of proteins is a major factor affecting amino acid usage in thermophiles^1, 3^. Similarly, a large body of work has demonstrated amino acid signatures associated with proteins from halophilic organisms^4–7^. The most common interpretation of these trends is that particular amino acid substitutions are selected through evolution to increase the stability and solubility of the folded conformation and enhance other structural properties such as flexibility^5^.

A complementary approach to interpreting patterns of amino acid composition is based on the energetics of amino acid synthesis. Energetic costs in terms of ATP requirements have been used to model protein expression levels in bacterial and yeast cells^8, 9^. Although ATP demands depend on environmental conditions^8^, a current limitation of ATP-based models is that they are derived for specific biosynthetic pathways, such as whether yeast are grown in respiratory or fermentative (i.e. aerobic or anaerobic) conditions^9^. A different class of models, based on thermodynamic analysis of the overall Gibbs energy of reactions to synthesize metabolites from inorganic precursors, quantifies the energetics of the reactions in terms of temperature, pressure, and chemical activities of all the species in the reactions, including those that define pH and oxidation-reduction potential^10^. Notably, the overall energetics for amino acid synthesis become more favorable, but to a different extent for each amino acid, between cold, oxidizing seawater and hot, reducing hydrothermal solution^11^. A recent systems biology study demonstrates tradeoffs between Gibbs energy of alternative pathways for amino acid synthesis and cofactor use efficiency (which affects ATP costs) in *E. coli* and suggests that pathway thermodynamics play a role in thermophilic adaptation^12^. Nevertheless, energetic models have not made much headway in relating metagenomic and geochemical data. This may be because few studies have asked whether specific changes in the chemical composition of biomolecules reflect specific environmental conditions.

To address this gap, here we claim that compositional analysis of proteins provides evidence for distinct adaptations to two types of environmental conditions: redox and salinity gradients. Because redox reactions are inherent in many aspects of metabolism, while hydration and dehydration reactions are essential for the synthesis of biomacromolecules^13^, our approach is shaped by the assumption that O_2_ and H_2_O are two primary components that link environmental conditions to the energetics of biomolecular synthesis. Thermodynamic considerations predict that redox gradients supply a driving force for changes in oxidation state of biomolecules (similar reasoning applies to oxygen content of proteins^14^), while salinity gradients, through the dehydrating potential associated with osmotic effects, exert a force that selectively alters the hydration state of biomolecules.

To test these predictions, we used two compositional metrics, the carbon oxidation state (*Z*_C_) and stoichiometric hydration state (*n*_H_2_O_). *Z*_C_ is computed from the chemical formulas of organic molecules, and takes values between the extremes of −4 (CH_4_) and +4 (CO_2_), although the range for particular classes of biomolecules is much smaller^15^. *n*_H_2_O_ is derived from the number of water molecules in theoretical formation reactions of proteins from basis species^16, 17^. Through analysis of representative metagenomic and metatranscriptomic datasets, we show that *Z*_C_ and *n*_H_2_O_ are most closely aligned with environmental redox and salinity gradients, respectively. These findings apply to freshwater and marine environments, but metagenomic and protein expression trends for hypersaline environments and halophiles deviate from the thermodynamic predictions, most likely due to optimizations of hydrophobicity and isoelectric point or specialized physiological adaptations.

In our previous study^18^, we compared one broad class of geochemical conditions (redox gradients) with one compositional metric for proteins (carbon oxidation state). Here, we expand the geobiochemical framework to two dimensions by considering another set of environments (salinity gradients) and another compositional metric (stoichiometric hydration state). A long-term research goal is to extend this framework to as many dimensions as there are thermodynamic components plus temperature and pressure. Further background on the concepts used in this paper, and their limitations, is provided below.

### Conceptual background

#### Intracellular conditions are maintained at physiological levels, so shouldn’t the comparisons be made with intracellular or local measurements of redox potential and salinity?

Available data show that physicochemical conditions in cells are not constant, but may be maintained in a narrower range relative to the environment. The most well studied example is for pH. In a compilation of data for different organisms, Figure 1 of ref. 19 shows that cytoplasmic pH varies from ca. 4.5 to 9.5 in external pH from 1 to 11.

**Figure 1.**
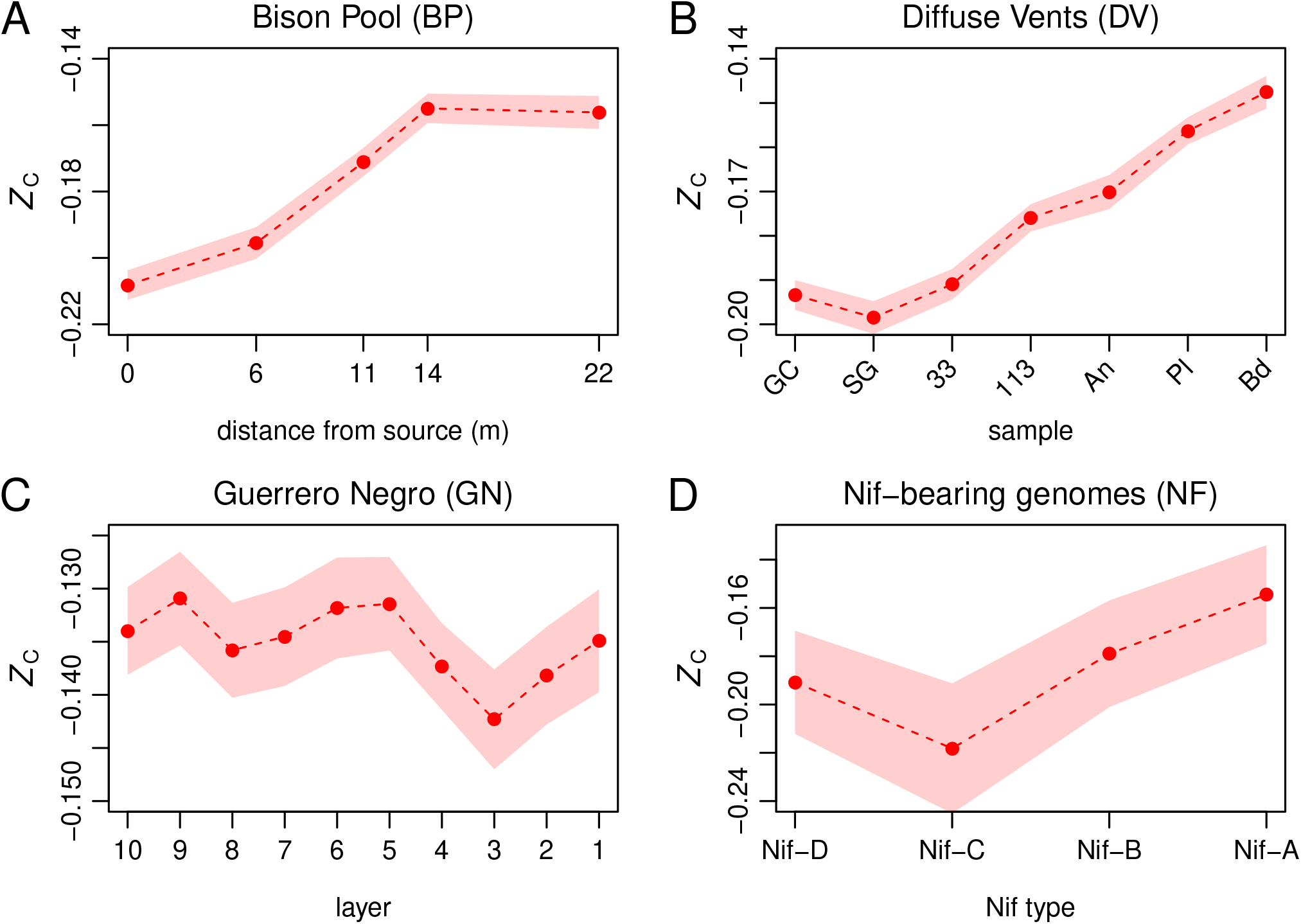
Changes in carbon oxidation state of proteins along redox gradients. Samples are ordered so that more oxidizing conditions are to the right. Filled circles indicate the average *Z*_C_ of protein sequences derived from metagenomes and the red shading denotes ±1 standard deviation (see Methods). Plots in (A)–(C) are adapted from Figure 2 of ref. 18. (A) At Bison Pool hot spring in Yellowstone National Park, the redox gradient is defined by increasing O_2_^40^ and sulfate and nitrate concentrations together with lower sulfide and ammonium^41^, and increasing oxidation-reduction potential (ORP)^31^. (B) For selected submarine diffuse vents, the most reducing conditions are found in ultramafic and mafic vent sites on the Mid-Cayman Rise with measured H_2_ concentrations of 11.3 mM at Ginger Castle (GC) and 0.41 mM at Shrimp Gulley #2 (SG)^36^. Lower hydrogen concentrations in diffuse vents on the Juan de Fuca Ridge, ranging from 0.1–1.5 μM at Marker 33 and Marker 113 to 12.5–27.7 μM at Anemone Vent^42^ (An), indicate that these fluids are less reducing. Marker 33 and 113 are considered to host stable methanogenic communities, whereas the relatively high H_2_ at Anemone is probably due to mixing in the shallow subsurface immediately before venting, lowering the availability of H_2_ for microbial utilization^42^, so An is placed to the right of Marker 33 and 113. The Anemone Plume (Pl) and background seawater (Bd) are both relatively oxidized environments. (C) The Guerrero Negro hypersaline microbial mat exhibits a daytime oxic-anoxic transition between ca. 1–2 mm and sulfidic conditions at greater depths^39^. Average sampling depths of layers 1 to 10 are 0.5, 1.5, 2.5, 3.5, 4.5, 5.5, 8, 16, 28, and 41.5 mm. (D) Values of *Z*_C_ calculated for proteomes of organisms bearing different homologs of nitrogenase. Nif-D is found exclusively in anaerobes, while Nif-A is present in some aerobic organisms^20^.

Cell membranes are permeable to uncharged species like hydrogen^19^, supporting the argument that oxidation-reduction conditions in the cytoplasm are affected by the external environment^20^. Oxygen diffuses rapidly through lipid membranes, depending on their composition and structure, and rates of diffusion increase with temperature^21^. Cell membranes are also permeable to water^22^. For *E. coli*, which grows most rapidly at about 0.3 OsM (osmolarity), increasing the extracellular osmotic strength from 0.1 to 1.0 OsM (approximately the osmotic concentration of seawater; BioNumbers^23^ BNID 100802) reduces the amount of free cytoplasmic water by more than half^22^. Halophiles, which thrive at even higher salinities, accumulate inorganic salts or organic solutes to maintain osmotic balance with the environment^6, 24^. This does not mean that intracellular hydration potential is constant. Intracellular water activity estimated from freezing point and cell composition data closely follows that of the growth medium, but is often offset to lower values^25^, perhaps due to macromolecular crowding effects^24^.

This brief review shows that the physicochemical conditions in cell interiors, at least under experimental conditions, are neither constant nor equal to the environment, but are sensitive to the environment. Ideally, we would like to compare the compositions of biomolecules to conditions actually measured inside cells or in the immediate surroundings of cells, but these measurements are generally not available for microbial communities in their natural environments, so we make comparisons with large-scale geochemical gradients, except for different layers of the Guerrero Negro microbial mat, where metagenomic and chemical data are available on the scale of millimeters.

#### The number of water molecules in formation reactions from basis species is not related to the water molecules released by the condensation of amino acids in protein synthesis. What does this number mean?

In the thermodynamics literature, a “formation reaction” represents the composition of a chemical species, either in terms of elements^26^, or in terms of other species^27^. When these other species are restricted in number to the minimum needed to represent the composition of all possible species in the system, they constitute a set of “basis species”, which can be thought of as the building blocks of the system, similar to the concept of thermodynamic components^28^. Therefore, a formation reaction from basis species is a mass-balanced, but non-unique, stoichiometric representation of the chemical composition of the protein. This type of reaction in general does not correspond to any biosynthetic mechanism, so to avoid confusion, we refer to these formation reactions as “theoretical formation reactions”; the number of water molecules in the theoretical formation reactions is the “stoichiometric hydration state”.

#### Proteins are synthesized by polymerization of L-alpha-amino acids, which is a condensation (dehydration) reaction. They are not synthesized directly from CO_2_, NH_3_, H_2_S, O_2_, and H_2_O (or a different set of basis species; see below), so how can hydration and oxidation in the environment affect the chemical composition of proteins?

The total metabolic demands for synthesizing proteins are not only for the polymerization reaction but also the formation of the amino acids themselves. To take one recent example, see the paper of Zhang *et al*.^29^, where the energy cost of proteins per amino acid in cancer cells was evaluated by averaging the contributions for amino acids making up the protein sequences. That paper refers to other papers that quantify the energetic cost for proteins using amino acid contributions in bacteria (*E. coli*) and yeast (*S. cerevisiae*)^8, 9^. In those papers, the energetic costs of proteins were computed per amino acid; likewise, in this paper, *Z*_C_ and *n*_H_2_O_ are defined as per-carbon and per-residue values, respectively, enabling comparisons between proteins of different length.

From a mechanistic standpoint, an analysis using any set of basis species is inadequate, since the number of basis species (five, corresponding to the number of elements, C, H, N, O, S) is smaller than the number of precursors and inorganic species that are actually involved in amino acid synthesis^12^. The use of O_2_, H_2_O, and other basis species to represent the composition of proteins reflects the hypothesis that they are meaningful physicochemical variables, even if they are not directly involved in the biosynthetic mechanisms for amino acids. The projection of amino acid composition (20-D) into the compositional space represented by basis species (5-D) can be thought of as a type of dimensionality reduction, but the variables are chosen based on a physicochemical hypothesis, unlike principal components analysis (PCA) or other unsupervised methods.

#### What are the evolutionary driving forces that may affect the chemical composition? Is it higher protein stability?

Thermodynamic models define the “cost” of a protein as a function of not only amino acid composition but also environmental conditions. Conceptually, this follows from Le Chatelier’s principle, in that increasing the chemical activity of a reactant (on the left-hand side of a reaction) drives the reaction toward the products, or in more general terms, that the overall Gibbs energy of a reaction depends on the activities of species in the reaction^10, 30^. Consider two proteins with different amino acid compositions, and therefore also different chemical compositions and theoretical formation reactions. The formation of the protein with more water as a reactant is theoretically favored by increasing the water activity, whereas the formation of the protein with more oxygen as a reactant is favored by increasing the oxygen activity. The water and oxygen activity are thermodynamic measures of hydration and oxidation potential and can be converted to other scales. Note that the number of O_2_ in theoretical formation reactions is closely related to the compositional metric used here, average oxidation state of carbon (see Methods).

This reasoning provides the theoretical justification for using chemical composition as an indicator for molecular adaptation to specific environmental conditions, but does not replace interpretations based on structural considerations. Halophilic organisms exhibit well-documented patterns of amino acid usage, including lower hydrophobicity and higher abundance of acidic residues, that impart greater stability, solubility, and flexibility of proteins^5^. These adaptations are reflected in lower values of the grand average of hydropathicity (GRAVY)^5, 7^ and/or isoelectric point of proteins (pI)^6^. In the Results, we compare the compositional analysis with GRAVY and pI and describe their different advantages.

#### It is well known that amino acid composition is affected by temperature and pH. In general, it is not controlled by a single environmental parameter. How can we tell if other environmental parameters affect oxidation and hydration state?

The redox gradients in hydrothermal systems are also temperature gradients (e.g. the mixing of seawater and hydrothermal fluid), and we have not attempted to disentangle the effects of temperature and redox conditions. However, our previous analysis of other redox gradients, including stratified hypersaline lakes, indicates that carbon oxidation state of biomolecules can vary even in systems where temperature changes are much smaller^18^. We claim that changes in oxidation state can be associated with one thermodynamic component of the system, and our goal in the present study is to explore the differences between this and one other component (represented by hydration state). Future work should also account for the effects of pH and temperature, which is possible using thermodynamic models for proteins^31^.

#### Is there an evolutionary or biosynthetic reason for choosing the QEC basis species?

The basis species used in this study for deriving the stoichiometric hydration state of proteins are glutamine, glutamic acid, cysteine, O_2_, and H_2_O (denoted QEC). The primary reason for choosing these basis species is to reduce the covariation between the metrics for oxidation and hydration state; that covariation is a mathematical consequence of projecting the elemental formulas of proteins into a particular compositional space, and may not reflect meaningful differences of chemical composition. There is nothing implied by the choice of basis species about evolutionary or biosynthetic mechanisms, and any set of basis species is thermodynamically valid, as long as they are the minimum number needed to represent the chemical composition of all the species in the system^28^. However, it is most convenient to select basis species that correspond to the controlling variables of the system. The QEC basis species has a biological rationale since glutamine and glutamic acid are often identified as highly abundant metabolites, and have been characterized as “nodal point” metabolites^32^. Other considerations are described in the first part of the Results.

## Results

### Choice of basis species

In this study we are concerned with the chemical formulas of primary amino acid sequences, not structural H_2_O molecules or the “hydration shell” of proteins. We aim to find a projection of the elemental composition of primary protein sequences (C, H, N, O, S) that clearly separates oxidation state and hydration state, which are represented by the stoichiometric numbers of O_2_ and H_2_O. Therefore, O_2_ and H_2_O are the only fixed requirements to have in the basis species. This leaves three basis species that must be able to be combined (along with O_2_ and H_2_O) in independent equations to give any number of C, N, and S.

There are no thermodynamic restrictions on the choice of basis species, but a biologically meaningful set is likely to comprise metabolites that have high network connectivity, i.e. are involved in reactions with many other metabolites. Reactions involving glutamine and glutamic acid, or its ionized form, glutamate, are major steps of nitrogen metabolism^33, 34^. Either methionine or cysteine would provide the required sulfur for the system, but cysteine is relevant as a constituent of the glutathione molecule, which has important roles in cellular redox chemistry^32^. These considerations support the proposal of the amino acids glutamine, glutamic acid, and cysteine (collectively abbreviated QEC) as a biologically relevant set of basis species for describing the chemical compositions of proteins^16^. These three amino acids are among the top eight amino acids ranked by number of reactions in a metabolic model for *Escherichia coli*^35^ (Glu: 52, Ser: 25, Asp: 23, Gln: 18, Ala: 15, Gly: 15, Met: 15, Cys: 13).

By reducing the covariation between *Z*_C_ and *n*_H_2_O_, the QEC basis species yields a more convenient projection than a common “default” choice of inorganic species, such as CO_2_, NH_3_, H_2_S, H_2_O, and O_2_, which commonly appear in overall catabolic reactions^30^ (see Methods for details). Furthermore, we used a residual correction (denoted “rQEC”) to reduce the remaining background correlation between *Z*_C_ and *n*_H_2_O_ (see Methods); this allows the identification of horizontal or vertical trends on *n*_H_2_O_–*Z*_C_ scatterplots to be associated with changes in only oxidation state or hydration state, respectively. The values of *n*_H_2_O_ for amino acids obtained with the rQEC derivation are listed in Table 1.

**Table 1.** Values of stoichiometric hydration state (*n*_H_2_O_) of amino acid residues from the rQEC derivation (see Methods). Standard one-letter abbreviations for the amino acids (AA) are used.

| AA | $n_{\text{H}_2\text{O}}$ | AA | $n_{\text{H}_2\text{O}}$ | AA | $n_{\text{H}_2\text{O}}$ | AA | $n_{\text{H}_2\text{O}}$ | AA | $n_{\text{H}_2\text{O}}$ |
| --- | --- | --- | --- | --- | --- | --- | --- | --- | --- |
| A | 0.369 | F | -2.568 | K | 0.763 | P | -0.354 | T | 0.569 |
| C | -0.025 | G | 0.478 | L | 0.660 | Q | -0.107 | V | 0.522 |
| D | -0.122 | H | -1.825 | M | 0.046 | R | 0.072 | W | -4.087 |
| E | -0.107 | I | 0.660 | N | -0.122 | S | 0.575 | Y | -2.499 |

### Comparison of redox and salinity gradients

To search for the hypothesized dehydration signal in metagenomic data, we began with redox gradients as a negative control. Submarine hydrothermal vents are zones of complex interactions between reduced endmember fluids and relatively oxidized seawater^36, 37^. Terrestrial hydrothermal systems, such as the hot springs in Yellowstone National Park, USA, provide a source of reduced fluids that are oxidized by degassing and mixing with air and surface groundwater as well as biological activity including sulfide oxidation^38^. Redox gradients can also develop at shorter length scales. The surface of the Guerrero Negro microbial mat (Baja California Sur, Mexico) is exposed to ~1 m deep hypersaline water that has ca. 200 μM dissolved oxygen, but in the mat, oxygen rises during the daytime and is depleted within a few millimeters, giving way to anoxic, then sulfidic conditions^39^.

Based on metagenomic data for these redox gradients^4, 40, 42–44^, Figure 1 shows that the carbon oxidation state of proteins increases dramatically in the outflow channel of Bison Pool (Fig. 1A) and between fluids from diffuse hydrothermal vents and relatively oxidizing seawater (Fig. 1B). It is noteworthy that intact polar lipids extracted from the microbial communities of Bison Pool and other alkaline hot springs also exhibit downstream increases in carbon oxidation state^41^. The *Z*_C_ of proteins increases more subtly toward the surface in the upper few mm of the Guerrero Negro microbial mat; it also increases at greater depths, perhaps due to heterotrophic degradation and/or horizontal gene transfer^18^ (Fig. 1C). Furthermore, an evolutionary trajectory associated with the occurrence of different homologs of nitrogenase (Nif) in anaerobic and aerobic organisms is characterized by increasing *Z*_C_ of the proteomes of these organisms^20^ (Fig. 1D).

With the exception of Guerrero Negro, these datasets exhibit larger changes in carbon oxidation state than stoichiometric hydration state (Fig. 2A). This is an expected outcome, as the redox gradients considered here do not have large changes in salinity. Concentrations of Cl^-^, a conservative ion, increase by less than 10% (6.1 to 6.6 mM) in the outflow of Bison Pool due to evaporation^40^. The diffuse vents considered here have concentrations of Cl^-^ between 515 and 624 mM, not greatly different from bottom seawater at 545 mM (Dataset S1 of ref. 36).

**Figure 2.**
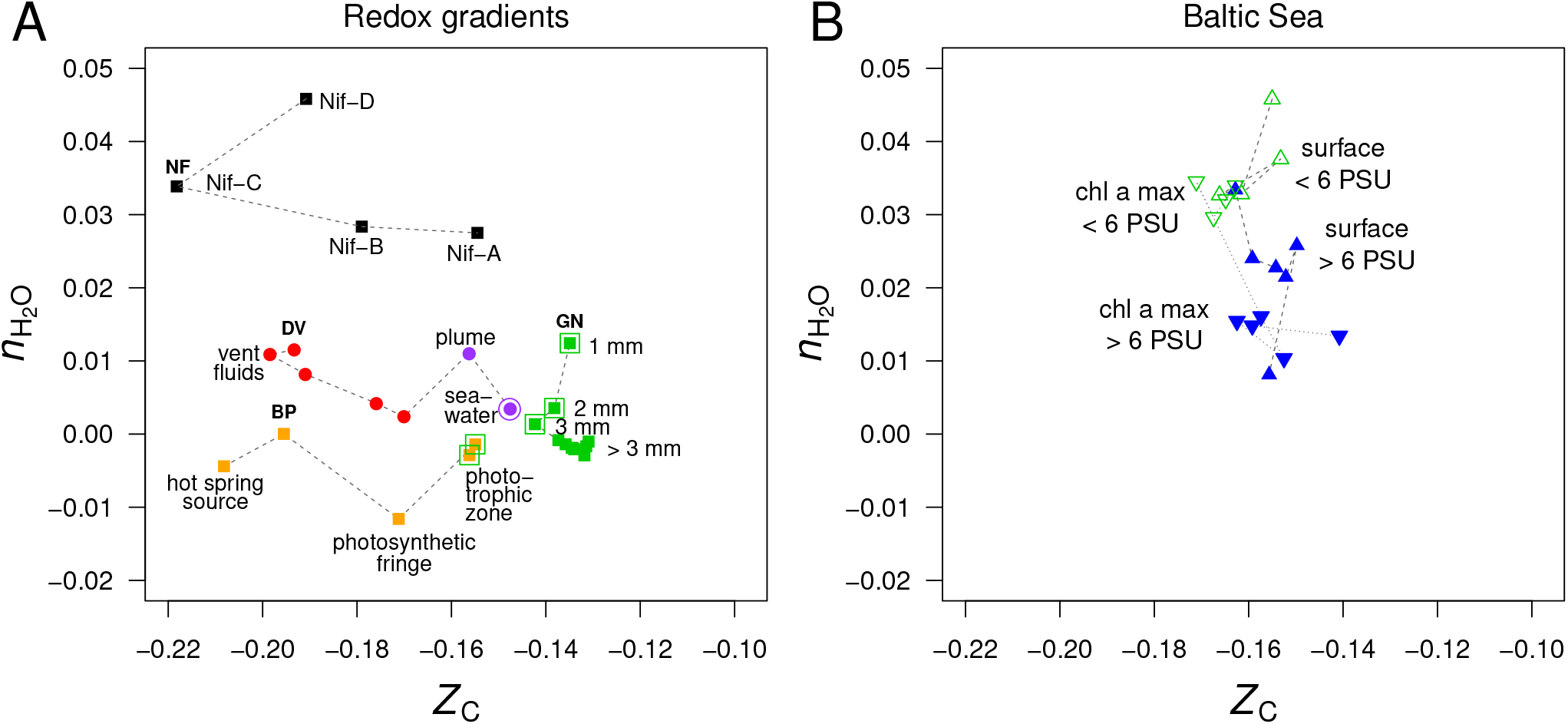
Compositional analysis of proteins in redox gradients and the Baltic Sea salinity gradient. (A) Redox gradients. Abbreviations and data sources are given in Fig. 1. Outlined symbols indicate samples in relatively oxidizing conditions. (B) Surface and deeper samples (chl a max: chlorophyll a maximum, 9–30 m deep) from the Baltic Sea transect. Metagenomes as described in ref. 45 were downloaded from iMicrobe^46^; data for the 0.1–0.8 μm size fraction are plotted here. Upward- and downward-pointing symbols, connected by dashed and dotted lines, represent surface and deeper samples, respectively, from stations along the transect at low salinity (< 6 PSU; open green symbols) and high salinity (> 6 PSU; filled blue symbols).

Turning our attention to a salinity gradient, the Baltic Sea exhibits a freshwater to marine transition over 1800 km, but dissolved oxygen at the surface is at or near saturation with air^45^, so this transect does not represent a redox gradient. For protein sequences derived from metagenomes in the 0.1–0.8 μm size fraction, we find large changes in stoichiometric hydration state along the Baltic Sea transect, but relatively small differences in the carbon oxidation state (Fig. 2B). This effect holds for samples from both the surface and chlorophyll a maximum (9–30 m deep).

### Multifactorial hydration effects

Metagenomic and metatranscriptomic data for different particle sizes are available for the Baltic Sea. The 0.1–0.8 μm and 0.8–3.0 μm size fractions represent free living bacteria, while the 3.0–200 μm fraction contains particle-associated bacteria with average larger genome sizes and greater inferred metabolic and regulatory capacity^45^. Fig. 3 shows that proteins derived from metagenomes of larger particles have lower *n*_H_2_O_ than those from the smallest size fraction. A plausible physical explanation is that the interiors of larger particles are sequestered to some extent from the surrounding aqueous environment, but evolutionary trends between unicellular and multicellular organisms could be another factor (see Discussion).

**Figure 3.**
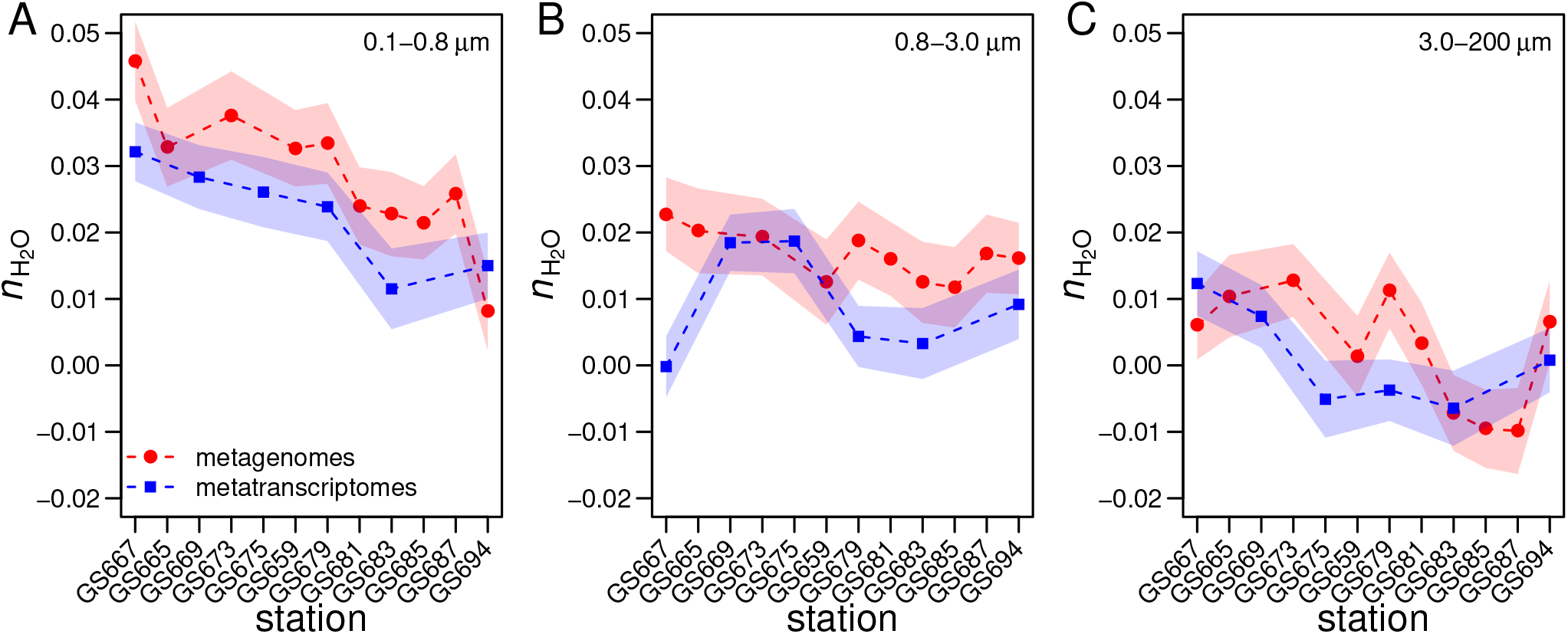
Variation of stoichiometric hydration state with increasing salinity from freshwater to marine conditions along the Baltic Sea transect (*x*-axis) and increasing particle size: (A) 0.1–0.8 μm, (B) 0.8–3.0 μm, (C) 3.0–200 μm. Sample IDs refer to the Sorcerer II Global Ocean Sampling Expedition; see ref. 45 for station descriptions. Protein sequences were derived from metagenomes^45^ (red circles) or metatranscriptomes^47^ (blue squares) of surface water samples. The freshwater metatranscriptome for the 0.8–3.0 μm sample from Torne Träsk (site GS667 in panel B) has an unusually low *n*_H_2_O_ and may be an outlier. Width of shading represents ±1 standard deviation in subsampled sequences (see Methods).

The Guerrero Negro microbial mat offers another opportunity to compare exposed and interior environments. Unlike *Z*_C_, which has a V-shaped pattern (Fig. 1C), *n*_H_2_O_ decreases throughout the mat, but the changes are most pronounced in the upper few millimeters (Fig. 2A). We speculate that this trend can be explained by greater exposure to bulk water at the surface of the mat or greater abundance of heterotrophic organisms with depth (see Discussion).

### Compositional trends in river and hypersaline environments

The Amazon river and ocean plume provide another example of a freshwater to marine transition, with salinities that range from below the scale of practical salinity units (PSU) in the river to 23-36 PSU in the plume^48, 49^. We used published metagenomic and metatranscriptomic data for filtered samples classified as free-living (0.2 to 2.0 μm) and particle-associated (2.0 to 156 μm)^48, 49^. River samples form a tight cluster on a plot of stoichiometric hydration state against carbon oxidation state of proteins; plume samples in the free-living size fraction are dispersed to lower *Z*_C_ whereas the particle-associated fraction shows very low values of *n*_H_2_O_ (Fig. 4A). Between the river and plume metatranscriptomes, *n*_H_2_O_ decreases noticeably but there is little overall difference in carbon oxidation state (Fig. 4B). The metatranscriptomes of particle-associated samples also have generally lower *n*_H_2_O_ than the free-living samples.

**Figure 4.**
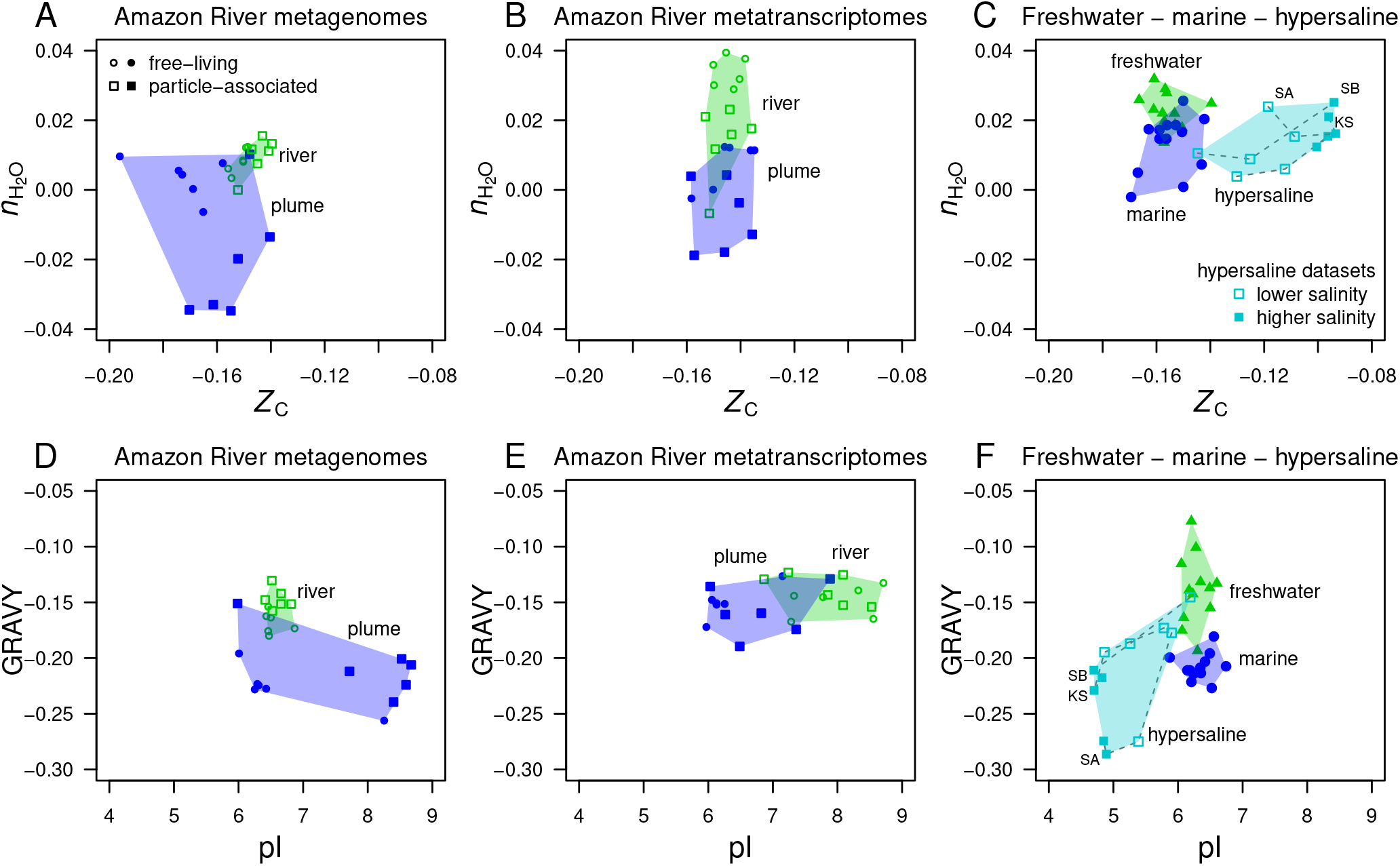
Compositional analysis and hydropathy and isoelectric point calculations for proteins from the Amazon River and plume and other metagenomes. Samples representing freshwater, marine, and hypersaline environments are indicated by the convex hulls colored green, blue, and light blue, respectively. (A) Metagenomic and (B) metatranscriptomic data for particle-associated and free-living fractions from the lower Amazon River^48^ (open symbols) and plume (filled symbols) in the Atlantic Ocean^49^. (C) Freshwater (lakes in Sweden and USA) and marine metagenomes considered in a previous comparative study^50^ and metagenomes from hypersaline environments including Kulunda Steppe soda lakes in Siberia, Russia^51^ (KS), Santa Pola salterns in Spain^52, 53^ (SA), and salterns in the South Bay of San Francisco, CA, USA^54^ (SB). Plots (D)–(F) show values of average hydropathicity (GRAVY) and isoelectric point (pI) of proteins for the same datasets.

Eiler *et al*.^50^ characterized microbial communities using metagenomic data for various freshwater samples (lakes in the USA and Sweden) and marine locations. Compositional analysis again reveals a relatively low *n*_H_2_O_ of proteins inferred from the marine metagenomes (Fig. 4C).

Hypersaline environments include evaporation ponds (salterns) and lakes in desert areas. The metagenomic data in our analysis come from the Santa Pola salterns in Spain^52, 53^, natural soda lakes of the Kulunda Steppe in Serbia^51^, and South Bay salterns in California, USA^54^. These hypersaline metagenomes have ranges of *n*_H_2_O_ of proteins that are similar to marine environments, but considerably higher *Z*_C_ (Fig. 4C). To interpret these results, we considered other factors that are known to influence the amino acid compositions of proteins in halophiles.

### Comparisons with hydropathy and isoelectric point

“Salt-in” halophilic organisms have proteins with relatively low isoelectric point that remain soluble in high salt concentration^52^. Notably, proteins with a lower pI also tend to have relatively high *Z*_C_ due to higher abundances of aspartic acid and glutamic acid, which are relatively oxidized (see refs. 11 and 55 and Fig. 6 in Methods). Consequently, the lower pI characteristic of “salt-in” organisms is also associated with an increase of carbon oxidation state (Fig. 4C). Because of the dominance of the pI shift, the increase of *Z*_C_ in this case is not an indicator of an environmental redox gradient.

Hydrophobic amino acids have high values on the hydropathicity scale (GRAVY) as well as relatively low values of *Z*_C_^55^. Therefore, values of GRAVY and *Z*_C_ for proteins are negatively correlated, but there is very little correlation between GRAVY and *n*_H_2_O_ (Supplementary Figure S1). Marine metagenomes have a lower GRAVY than most of the freshwater samples, and hypersaline metagenomes are shifted to both lower GRAVY and pI (Fig. 4F). However, there are irregular trends in the Amazon River data. Compared to the river, the plume metagenomes exhibit lower GRAVY and either higher or lower pI (Fig. 4D). Similarly, other authors have reported that although lower pI is a signature of many hypersaline environments, it does not distinguish marine from low-salinity environments^56^. On the other hand, the plume metatranscriptomes do show decreased pI but no discernible trend in GRAVY (Fig. 4E).

Considering all the variables and datasets shown in Fig. 4, only *n*_H_2_O_ exhibits a consistent trend – toward lower values – in marine compared to freshwater samples. However, the decrease of stoichiometric hydration state between freshwater and marine environments does not continue into hypersaline environments.

### Compositional analysis of differentially expressed proteins

To probe into the physiological basis for the patterns found in metagenomes and metatranscriptomes, we considered the changes in transcripts and protein expression levels when cells are exposed to hyperosmotic conditions in the laboratory. Differences in the mean values of *n*_H_2_O_ and *Z*_C_ for differentially expressed proteins reported in 13 studies on non-halophilic bacterial and eukaryotic cells were summarized in a previous publication^17^. We augmented that compilation with data from an additional seven studies on bacteria and archaea^57–63^, including four halophiles (Table 2). The reported significantly differentially expressed genes or proteins were mapped to UniProt accession numbers^66^; proteins with unavailable UniProt IDs, those with ambiguous expression (appearing in both the down- and up-regulated groups), and duplicates were removed.

**Table 2.**
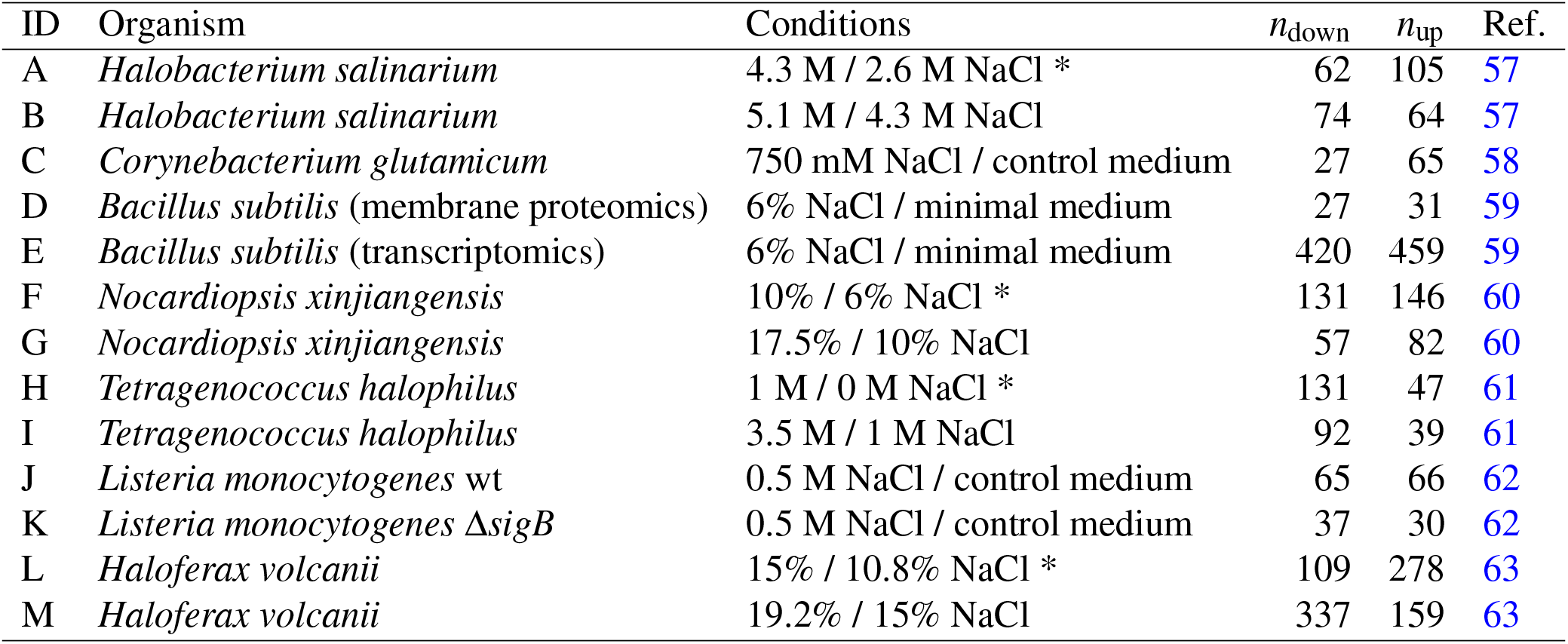
Organisms, growth conditions, number of differentially expressed proteins, and sources of data for hyperosmotic and hypoosmotic stress experiments. All data are for proteomics experiments unless noted otherwise. Hypoosmotic experiments, where the control condition has lower than optimal salinity, are marked with *. Units for NaCl concentrations are taken from the references; approximate conversions between molarity and weight percent are 1 M NaCl ≈ 6%, 2.5 M NaCl ≈ 13%, 4 M NaCl ≈ 20%. Data sources: (A, B) Tables 1 and 2 of ref. 57. (C) Supplementary Table 8 of ref. 58. Only proteins with consistent expression ratios (all > 1 or all < 1) at each time point (15, 60, and 180 min.) were included. (D, E) Supplementary Tables 2 (membrane proteomics) and 3 (transcriptomics) of ref. 59. The cytosol proteomics experiment was not included because of the low number of proteins identified with a significant expression difference. Only proteins with consistent expression ratios in at least 3 of 4 time points (10, 30, 60, and 120 min.) were included. (F, G) Table S-1 of ref. 60. Values of reporter intensities at each condition (6%, 10%, and 17.5% NaCl) were quantile normalized and used to compute intensity ratios (10% / 6% NaCl and 17.5% / 10% NaCl). Only proteins with expression ratios > 1.3 in either direction^60^, *p*-values < 0.05, and at least 2 peptides were included. (H, I) Tables S2 and S3 of ref. 61. (J, K) Tables S1–S6 of ref. 62. For each of the wild-type and Δ*sigB* mutant, only proteins that were identified in extracellular vesicles in a single condition (0.5 M salt stress or without salt stress) were included. (L, M) Supporting Table 1C of ref. 63. Only proteins with at least 2-fold expression difference and marked as significant were included.

| ID | Organism | Conditions | $n_{\text{down}}$ | $n_{\text{up}}$ | Ref. |
| --- | --- | --- | --- | --- | --- |
| A | *Halobacterium salinarium* | 4.3 M / 2.6 M NaCl * | 62 | 105 | 57 |
| B | *Halobacterium salinarium* | 5.1 M / 4.3 M NaCl | 74 | 64 | 57 |
| C | *Corynebacterium glutamicum* | 750 mM NaCl / control medium | 27 | 65 | 58 |
| D | *Bacillus subtilis* (membrane proteomics) | 6% NaCl / minimal medium | 27 | 31 | 59 |
| E | *Bacillus subtilis* (transcriptomics) | 6% NaCl / minimal medium | 420 | 459 | 59 |
| F | *Nocardiopsis xinjiangensis* | 10% / 6% NaCl * | 131 | 146 | 60 |
| G | *Nocardiopsis xinjiangensis* | 17.5% / 10% NaCl | 57 | 82 | 60 |
| H | *Tetragenococcus halophilus* | 1 M / 0 M NaCl * | 131 | 47 | 61 |
| I | *Tetragenococcus halophilus* | 3.5 M / 1 M NaCl | 92 | 39 | 61 |
| J | *Listeria monocytogenes* wt | 0.5 M NaCl / control medium | 65 | 66 | 62 |
| K | *Listeria monocytogenes* $\Delta\text{sigB}$ | 0.5 M NaCl / control medium | 37 | 30 | 62 |
| L | *Haloferax volcanii* | 15% / 10.8% NaCl * | 109 | 278 | 63 |
| M | *Haloferax volcanii* | 19.2% / 15% NaCl | 337 | 159 | 63 |

The combined data are plotted in Fig. 5A. A lower *n*_H_2_O_ of up-regulated proteins characterizes most of the hyperosmotic stress experiments, including microbial proteomes and transcriptomes and eukaryotic proteomes. Notably, the datasets cluster near a difference in *n*_H_2_O_ of ca. −0.02, which is similar to the differences between freshwater and marine metagenomes and metatranscriptomes described above. An opposite pattern emerges for increasing salinity from *hypoosmotic* conditions to optimal growth salinity for some halophilic organisms, indicated by large squares in Fig. 5. These datasets represent the archaeon *Halobacterium salinarium* and bacteria *Nocardiopsis xinjiangensis* and *Tetragenococcus halophilus* (points labeled A, F, and H). Interestingly, datasets for *hyperosmotic* stress from the same studies show lower Δ*n*_H_2_O_ (negative values in two of three datasets; points labeled B, G, I), which is closer to the general pattern for hyperosmotic stress experiments in non-halophilic organisms.

**Figure 5.**
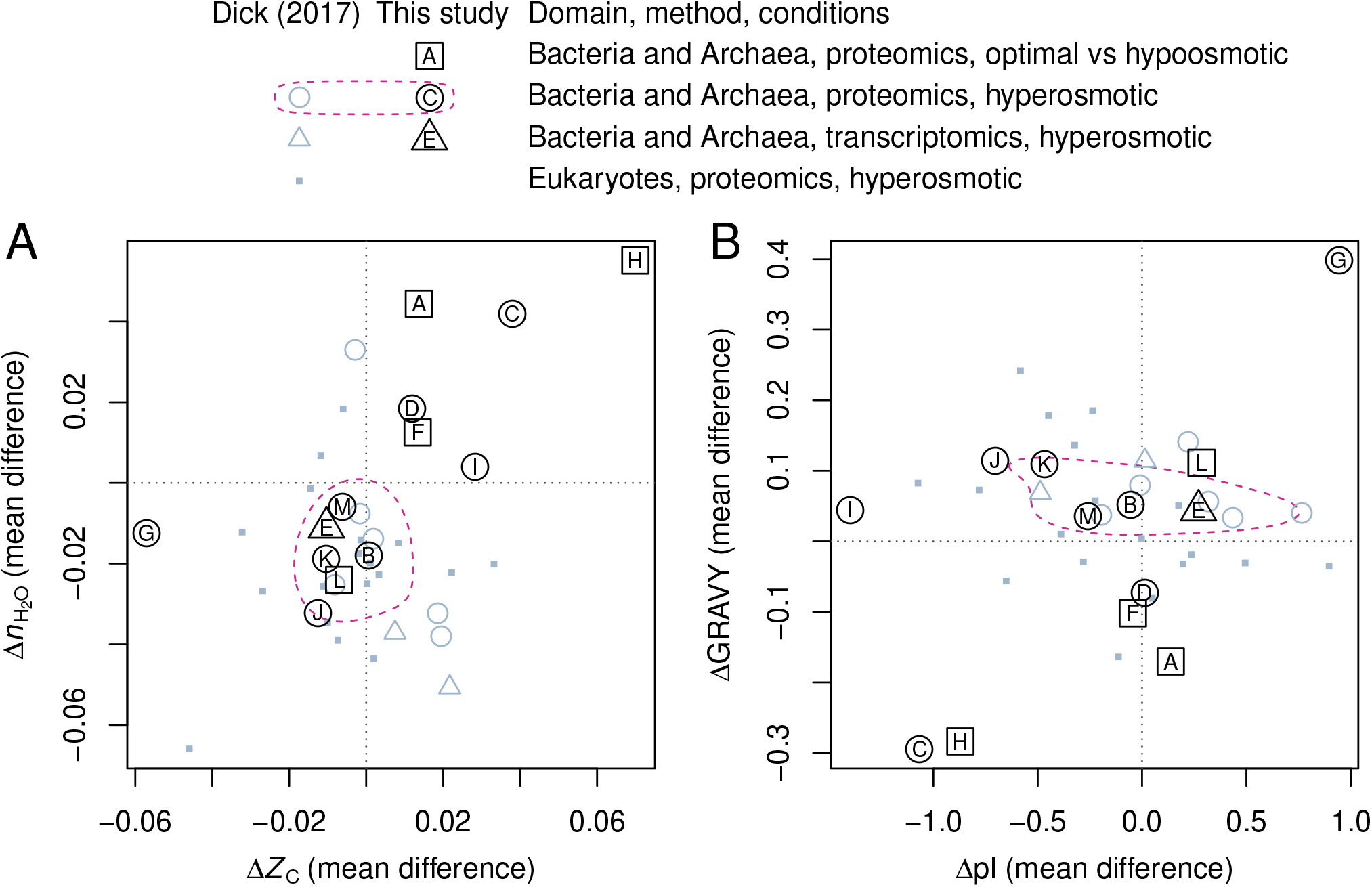
Compositional analysis of differentially expressed proteins in osmotic stress experiments. (A) Mean differences of *n*_H_2_O_ and *Z*_C_ between up- and down-regulated proteins in response to increasing osmotic strength. Gray symbols represent data compiled by Dick (2017) (ref. 17). Black symbols labeled with letters represent data added in this study (Table 2). The dashed outline indicates the 50% credible region for highest probability density for proteomics data for Bacteria and Archaea under hyperosmotic conditions (code adapted from the HPDregionplot function in R package emdbook^64^ with kernel density estimates that were computed with function kde2d in R package MASS^65^). (B) Mean differences of GRAVY and pI for the same datasets.

The mean value of GRAVY increases for differentially expressed proteins in most hyperosmotic stress experiments (Fig. 5B). This trend is opposite that found for halophilic adaptation^5^ and the metagenomic comparisons described above. We conclude that *n*_H_2_O_ is a more generally applicable metric, since it records decreasing hydration state along salinity gradients in the Baltic Sea and Amazon River, between other freshwater and marine metagenomes, and in protein expression under hyperosmotic stress.

## Discussion

Based on mass-action effects in thermodynamics, we predicted that carbon oxidation state of proteins (*Z*_C_) should increase toward more oxidizing conditions and that stoichiometric hydration state (*n*_H_2_O_) should decrease toward higher salinity. We found that proteins inferred from metagenomes in redox gradients associated with submarine hydrothermal systems and a Yellowstone hot spring exhibit large changes of *Z*_C_, whereas regional salinity gradients, including the Baltic Sea freshwater-marine transect and Amazon River and plume, are characterized by large changes of *n*_H_2_O_. Although *n*_H_2_O_ decreases between freshwater and marine environments, the trend does not continue into hypersaline environments.

Laboratory experiments provide another view of the effects of osmotic conditions on protein expression. Compositional analysis of proteomic data indicates a net loss of H_2_O from differentially expressed proteins in many non-halophilic organisms in hyperosmotic stress experiments. Differentially expressed proteins in halophiles show a more complex response: positive Δ*n*_H_2_O_ between hypoosmotic conditions and optimal growth salinities, and lower (sometimes negative) Δ*n*_H_2_O_ between optimal and hyperosmotic conditions. We speculate that the net uptake or release of water from proteomes in response to changing growth conditions could affect cell metabolism, perhaps contributing to the effects of metabolically produced water on microbial growth at low water activities^67^.

Limited accessibility to the aqueous phase might play a role for the lower *n*_H_2_O_ in particles compared to free-living fractions in the Baltic Sea and Amazon River and in the deeper layers of the Guerrero Negro microbial mat. However, it should be noted that larger particles are often associated with more multicellular and eukaryotic assemblages^68^. A lower average *n*_H_2_O_ is also apparent for human proteins (mean defined to be zero) compared to *E. coli* (see Methods) and many of the metagenomic and metatranscriptomic datasets considered here, as indicated by mostly positive values of *n*_H_2_O_ in Figs. 2–4. Similarly, cancer tissues are characterized by up-regulation of unicellular genes^69^ as well as proteins with higher *n*_H_2_O_ compared to normal tissue^17^. More work is needed to confirm whether hydration state decreases through the evolution of multicellularity, as suggested by these preliminary observations.

Another important evolutionary transition is the emergence of heterotrophic metabolism, which is considered to be a later innovation than autotrophic core metabolism^13, 33^. It is notable that the deeper layers of the Guerrero Negro mat show greater evidence for heterotrophic metabolism^4^; likewise, heterotrophs at the “photosynthetic fringe” in Bison Pool may outcompete the autotrophs that dominate at higher and lower temperatures^40^. These putative heterotroph-rich zones show locally lower values of *n*_H_2_O_ (Fig. 2A). If decreasing stoichiometric hydration state is a common theme across evolutionary transitions, then the relatively high *n*_H_2_O_ in the proteomes of organisms carrying the ancestral nitrogenase Nif-D^20^ (Fig. 2A) is not unexpected. The findings of this study underscore an opportunity for integrating hydration state into evolutionary models that already consider changes in oxidation state or oxygen content of proteins^14, 20^.

## Methods

### Prediction of protein sequences

Protein sequences were predicted from metagenomic reads using a previously described workflow^18^. Briefly, reads were trimmed, filtered, and dereplicated using scripts adapted from the MG-RAST pipeline^70^. For metatranscriptomic datasets, ribosomal RNA sequences were removed using SortMeRNA^71^. Protein-coding sequences were identified using FragGeneScan^72^, and the amino acid sequences of the predicted proteins were used in further calculations. For large datasets, only a portion of the available reads was processed (at least 500,000 reads; see Supplementary Table S1). This reduces the computational requirements without noticeably affecting the calculated average compositions^18^.

### Average oxidation state and stoichiometric hydration state

The average oxidation state of carbon (*Z*_C_) measures the degree of oxidation of carbon atoms in organic molecules. For a protein with chemical formula C_*c*_H_*h*_N_*n*_O_*o*_S_*s*_, the value of *Z*_C_ can be calculated from^31, 55^

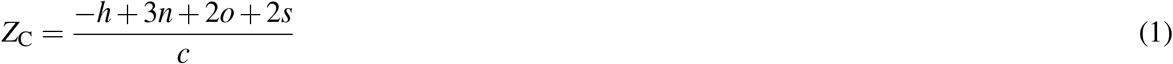

Using the basis species CO_2_, NH_3_, H_2_S, H_2_O, and O_2_ (referred to here as CHNOS), the theoretical formation reaction of an amino acid, for example alanine (C_3_H_7_NO_2_), is

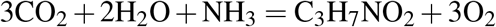

and the oxygen and water content of alanine (*n*_O_2__ = −3 and *n*_H_2_O_ = 2) are the opposite of the coefficients on O_2_ and H_2_O in the reaction. Stoichiometric coefficients for amino acids calculated using the CHNOS basis were used to make Fig. 6A–B.

**Figure 6.**
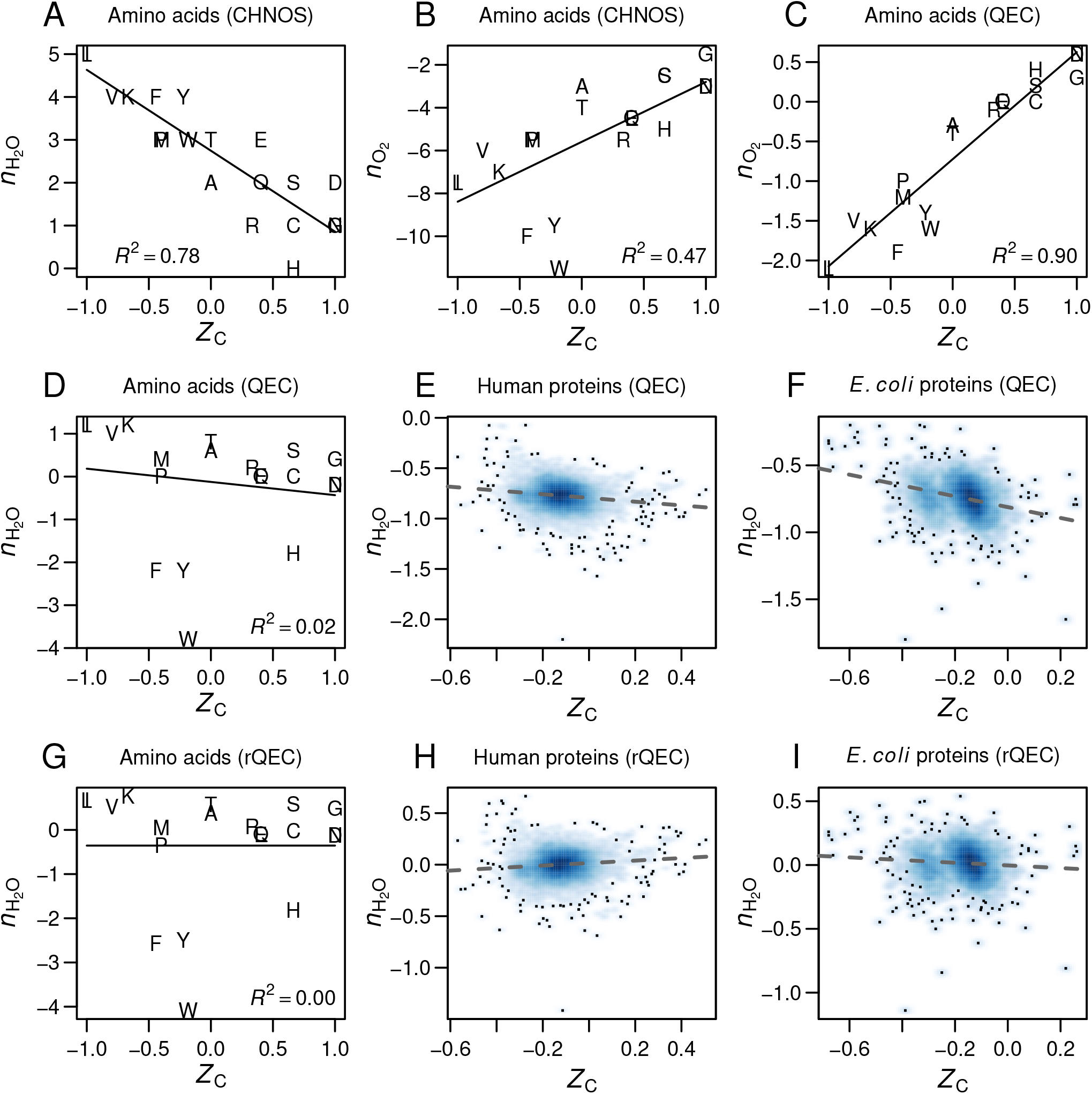
Comparison of compositional metrics for oxidation state and hydration state with different sets of basis species (CHNOS and QEC) and derivation of the residual correction (rQEC). (A, B) Number of H_2_O and O_2_ in the theoretical formation reactions of amino acids from CO_2_–NH_3_–H_2_S–H_2_O–O_2_ (CHNOS) are plotted against carbon oxidation state (*Z*_C_), which is also computed from the chemical formula but does not depend on the choice of basis species. Linear models and *R*^2^ values were calculated using the lm function in R^73^. (C, D) Changing the basis species to glutamine–glutamic acid–cysteine–H_2_O–O_2_ (QEC) strengthens the association between *Z*_C_ and *n*_O_2__ and decreases that between *Z*_C_ and *n*_H_2_O_. However, there is still a noticeable negative correlation between *Z*_C_ and *n*_H_2_O_, which is also visible in scatterplots of all proteins in (E) *H. sapiens* and (F) *E. coli* K12 (UniProt^66^ reference proteomes UP000005640 and UP000000625). (G) Residuals from the linear model in (D) minus a constant of 0.355 yield values for the stoichiometric hydration state (rQEC) of amino acids. (H, I) Stoichiometric hydration states of proteins calculated from the rQEC values. The constant was defined so that the mean *n*_H_2_O_ for human proteins equals zero.

Using glutamine (C_5_H_10_N_2_O_3_), glutamic acid (C_5_H_9_NO_4_), cysteine (C_3_H_7_NO_2_S), H_2_O, and O_2_ (the QEC basis species), the theoretical formation reaction of alanine is

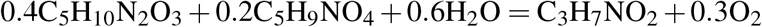

showing that the oxygen and water content are *n*_O_2__ = −0.3 and *n*_H_2_O_ = 0.6. The QEC basis was used to make Fig. 6C–F.

The compositions of amino acids in the CHNOS basis exhibit a strong negative correlation between *Z*_C_ and *n*_H_2_O_ (Fig. 6A), but a relatively weak correlation between *Z*_C_ and *n*_O_2__ (Fig. 6B). The QEC basis provides a much stronger association between *Z*_C_ and *n*_O_2__ (which can both be interpreted as metrics for oxidation state) and also greatly reduces the coupling between *Z*_C_ and *n*_H_2_O_ (Fig. 6C–D). However, there is a small remaining negative correlation for amino acids (Fig. 6D), which is also visible in whole-proteome data for humans and *E. coli* (Fig. 6E–F). We derived values of *n*_H_2_O_ by taking the residuals of the linear model for amino acids (Fig. 6D), then subtracting a constant so that the mean for all human proteins equals zero. This derivation, which we refer to as “rQEC”, gives the residual-corrected stoichiometric hydration state for each amino acid, which is plotted in Fig. 6G and listed in Table 1. Even with the residual correction for amino acids, there remain slightly positive and negative correlations for human and *E. coli* proteins (Fig. 6H–I). As noted above, the mean *n*_H_2_O_ for human proteins was defined to be zero; the mean for proteins in *E. coli* is somewhat greater, at 0.014.

### Compositional metrics for proteins and metagenomes

The stoichiometric hydration state of proteins was calculated by multiplying the values from Table 1 by the frequencies of the amino acids and dividing the sum by the number of amino acids. The average oxidation state of carbon was also calculated from the amino acid values (see Table 1 of ref. 31) and frequencies. Unlike *n*_H_2_O_, averages for *Z*_C_ must be weighted by the number of carbon atoms in each amino acid. For example, *Z*_C_ of the dipeptide Ala-Gly can be calculated as (3 × 0 + 2 × 1) / (3 + 2), where 3 and 2 are the numbers of carbon atoms and 0 and 1 are the *Z*_C_ of Ala and Gly, respectively. This result (0.4) can be checked by combining Eq. 1 with the chemical formula of alanylglycine (C_5_H_10_N_2_O_3_).

Means and standard deviations of *Z*_C_ and *n*_H_2_O_ were calculated for 100 random subsamples of protein sequences from each metagenomic or metatranscriptomic dataset. The numbers of sequences included in the subsamples were chosen to give a total length closest to 50,000 amino acids on average.

### Amino acid composition of proteomes of Nif-bearing organisms

Amino acid compositions of all proteins for each bacterial, archaeal, and viral taxon in the NCBI Reference Sequence (RefSeq) database^74^ were compiled from RefSeq release 95 (July 2019). Scripts to do this, and the resulting data file of amino acid compositions of 36,425 taxa, are available in version 1.0.1 of the JMDplots package (see Data Availability). Names of organisms containing different nitrogenase (Nif) homologs were extracted from Supplemental Table 1A of ref. 20. These names were matched to the closest organism name in RefSeq. Duplicated species (represented by different strains) were removed, as were matching organisms with fewer than 1000 RefSeq protein sequences. As a result, the numbers of organisms included in these calculations (Nif-A: 157, Nif-B: 69, Nif-C: 14, Nif-D: 7) are less than those identified in ref. 20. Note that values of *Z*_C_ calculated here (Fig. 1D) are lower than those shown in Fig. 5 of ref. 20. This difference is associated with the weighting by carbon number (described above), which was not done in ref. 20.

### GRAVY and pI

The grand average of hydropathicity (GRAVY) was calculated using published hydropathy values for amino acids^75^. The isoelectric point was calculated using published p*K* values for terminal groups^76^ and sidechains^77^; however, the calculation does not implement position-specific adjustments for some groups^77^. The charge for each ionizable group was precalculated from pH 0 to 14 at intervals of 0.01, and the isoelectric point was computed as the pH where the sum of charges of all groups in the protein is closest to zero. These calculations were implemented as new functions in version 0.1.5 of the canprot package^17^ for R^73^ (see Data Availability). Comparisons for a few proteins show that the calculated values of GRAVY and pI are equal to those obtained with the ProtParam tool^78^. Means of GRAVY and pI were calculated for subsampled sequences as described above.

## Data Availability

All metagenomic and metatranscriptomic data analyzed here were obtained from public databases using the accession numbers listed in Supplementary Table S1 or identified previously for redox gradients (Supplementary Material of ref. 18). The data used for Fig. 5 are available in version 0.1.5 of the canprot R package (https://github.com/jedick/canprot), which is deposited at Zenodo (https://doi.org/10.5281/zenodo.3676560). Data used to make the remaining figures, including amino acid compositions of subsampled sequences from the metagenomic and metatranscriptomic data, are available in version 1.0.1 of the JMDplots R package (https://github.com/jedick/JMDplots), which is deposited at Zenodo (https://doi.org/10.5281/zenodo.3676632). The JMDplots package also has the code used to make all the figures in this paper; see the “gradH2O” vignette in the package.

## Acknowledgements

Thanks to Saroj Poudel for helpful comments on an earlier version of the manuscript. This work was supported by funding from the State Key Laboratory of Organic Geochemistry (Grant No. SKLOG-201928 to J.D.).

## Supplementary Material

**Figure S1.**
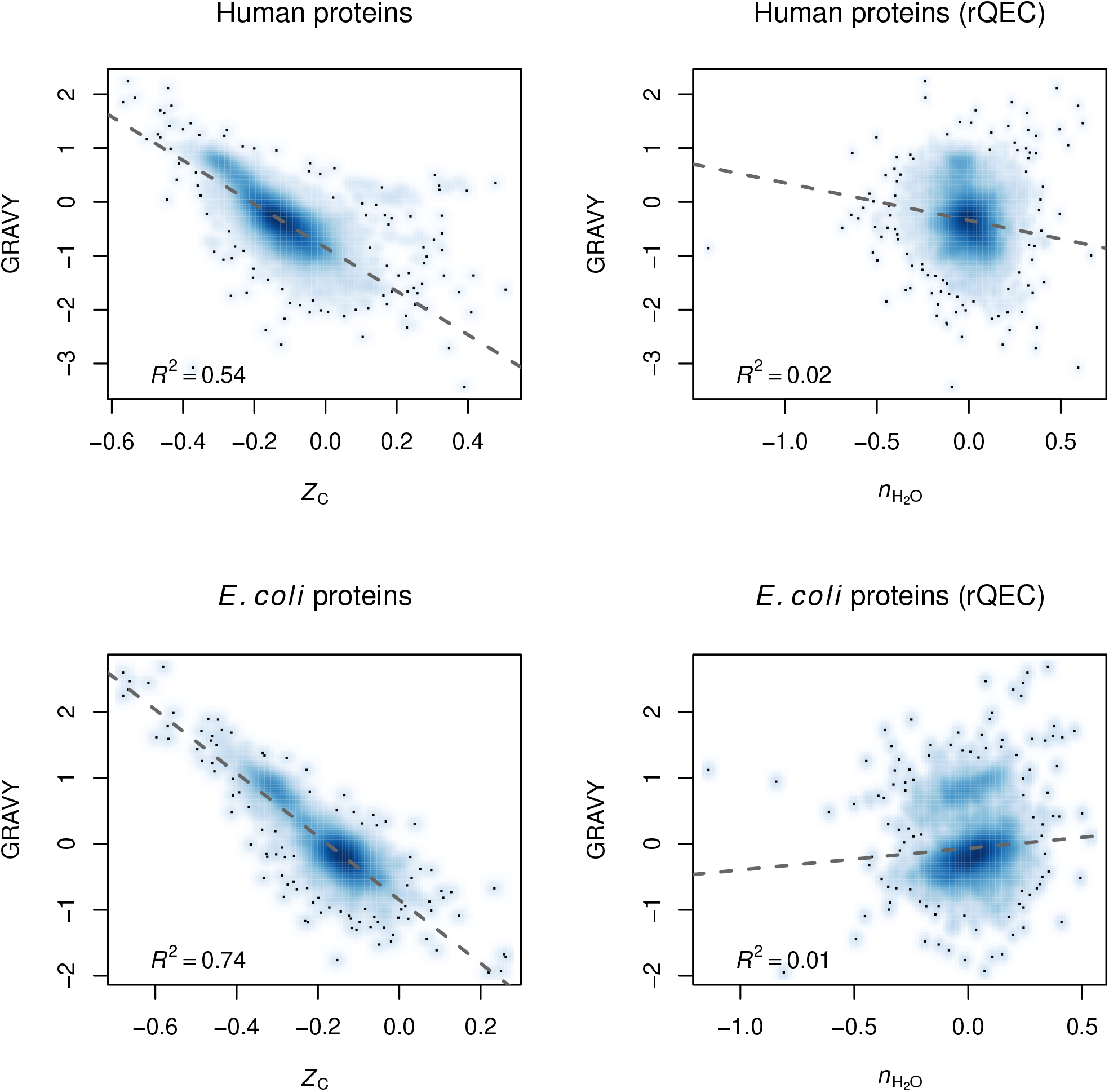
Scatterplots of GRAVY vs *Z*_C_ and *n*_H_2_O_. Proteins in *H. sapiens* and *E. coli* were downloaded from UniProt, and GRAVY, *Z*_C_, and *n*_H_2_O_ were calculated as described in the Methods. Linear models and *R*^2^ values were calculated using the lm function in R.

## References

1. Zeldovich, K. B., Berezovsky, I. N. & Shakhnovich, E. I. Protein and DNA sequence determinants of thermophilic adaptation. PLoS Comput. Biol. 3, 62–72, DOI: 10.1371/journal.pcbi.0030005 (2007).

2. Alsop, E. B., Boyd, E. S. & Raymond, J. Merging metagenomics and geochemistry reveals environmental controls on biological diversity and evolution. BMC Ecol. 14, 16, DOI: 10.1186/1472-6785-14-16 (2014).

3. Sterner, R. & Liebl, W. Thermophilic adaptation of proteins. Critical Rev. Biochem. Mol. Biol. 36, 39–106, DOI: 10.1080/20014091074174 (2001).

4. Kunin, V. et al. Millimeter-scale genetic gradients and community-level molecular convergence in a hypersaline microbial mat. Mol. Syst. Biol. 4, 198, DOI: 10.1038/msb.2008.35 (2008).

5. Paul, S., Bag, S. K., Das, S., Harvill, E. T. & Dutta, C. Molecular signature of hypersaline adaptation: insights from genome and proteome composition of halophilic prokaryotes. Genome Biol. 9, R70, DOI: 10.1186/gb-2008-9-4-r70 (2008).

6. Oren, A. Life at high salt concentrations, intracellular KCl concentrations, and acidic proteomes. Front. Microbiol. 4, 315, DOI: 10.3389/fmicb.2013.00315 (2013).

7. Boyd, E. S. et al. [FeFe]-hydrogenase abundance and diversity along a vertical redox gradient in Great Salt Lake, USA. Int. J. Mol. Sci. 15, 21947–21966, DOI: 10.3390/ijms151221947 (2014).

8. Akashi, H. & Gojobori, T. Metabolic efficiency and amino acid composition in the proteomes of *Escherichia coli* and *Bacillus subtilis*. Proc. Natl. Acad. Sci. 99, 3695–3700, DOI: 10.1073/pnas.062526999 (2002).

9. Wagner, A. Energy constraints on the evolution of gene expression. Mol. Biol. Evol. 22, 1365–1374, DOI: 10.1093/molbev/msi126 (2005).

10. Shock, E. L. et al. Quantifying inorganic sources of geochemical energy in hydrothermal ecosystems, Yellowstone National Park, USA. Geochim. Cosmochim. Acta 74, 4005–4043, DOI: 10.1016/j.gca.2009.08.036 (2010).

11. Amend, J. P. & Shock, E. L. Energetics of amino acid synthesis in hydrothermal ecosystems. Science 281, 1659–1662, DOI: 10.1126/science.281.5383.1659 (1998).

12. Du, B., Zielinski, D. C., Monk, J. M. & Palsson, B. O. Thermodynamic favorability and pathway yield as evolutionary tradeoffs in biosynthetic pathway choice. Proc. Natl. Acad. Sci. 115, 11339–11344, DOI: 10.1073/pnas.1805367115 (2018).

13. Braakman, R. & Smith, E. The compositional and evolutionary logic of metabolism. Phys. Biol. 10, 011001, DOI: 10.1088/1478-3975/10/1/011001 (2013).

14. Acquisti, C., Kleffe, J. & Collins, S. Oxygen content of transmembrane proteins over macroevolutionary time scales. Nature 445, 47–52, DOI: 10.1038/nature05450 (2007).

15. Amend, J. P., LaRowe, D. E., McCollom, T. M. & Shock, E. L. The energetics of organic synthesis inside and outside the cell. Philos. Transactions Royal Soc. B: Biol. Sci. 368, 20120255, DOI: 10.1098/rstb.2012.0255 (2013).

16. Dick, J. M. Proteomic indicators of oxidation and hydration state in colorectal cancer. PeerJ 4, e2238, DOI: 10.7717/peerj.2238 (2016).

17. Dick, J. M. Chemical composition and the potential for proteomic transformation in cancer, hypoxia, and hyperosmotic stress. PeerJ 5, e3421, DOI: 10.7717/peerj.3421 (2017).

18. Dick, J. M., Yu, M., Tan, J. & Lu, A. Changes in carbon oxidation state of metagenomes along geochemical redox gradients. Front. Microbiol. 10, 120, DOI: 10.3389/fmicb.2019.00120 (2019).

19. Slonczewski, J. L., Fujisawa, M., Dopson, M. & Krulwich, T. A. Cytoplasmic pH measurement and homeostasis in bacteria and archaea. In Poole, R. K. (ed.) Advances in Microbial Physiology, vol. 55, 1–79, DOI: 10.1016/S0065-2911(09)05501-5 (Academic Press, New York, 2009).

20. Poudel, S. et al. Electron transfer to nitrogenase in different genomic and metabolic backgrounds. J. Bacteriol. 200, e00757–17, DOI: 10.1128/JB.00757-17 (2018).

21. Möller, M. N. et al. Solubility and diffusion of oxygen in phospholipid membranes. Biochimica Biophys. Acta (BBA) - Biomembr. 1858, 2923–2930, DOI: 10.1016/j.bbamem.2016.09.003 (2016).

22. Record, M. T., Jr., Courtenay, E. S., Cayley, D. S. & Guttman, H. J. Responses of *E. coli* to osmotic stress: large changes in amounts of cytoplasmic solutes and water. Trends Biochem. Sci. 23, 143–148, DOI: 10.1016/S0968-0004(98)01196-7 (1998).

23. Milo, R., Jorgensen, P., Moran, U., Weber, G. & Springer, M. BioNumbers—the database of key numbers in molecular and cell biology. Nucleic Acids Res. 38, D750–D753, DOI: 10.1093/nar/gkp889 (2010).

24. Garner, M. M. & Burg, M. B. Macromolecular crowding and confinement in cells exposed to hypertonicity. Am. J. Physiol. 266, C877–C892, DOI: 10.1152/ajpcell.1994.266.4.C877 (1994).

25. Chirife, J., Fontan, C. F. & Scorza, O. C. The intracellular water activity of bacteria in relation to the water activity of the growth medium. J. Appl. Bacteriol. 50, 475–479, DOI: 10.1111/j.1365-2672.1981.tb04250.x (1981).

26. Warn, J. R. W. & Peters, A. P. H. Concise Chemical Thermodynamics (CRC Press, 1996), 2nd edn.

27. May, P. M. & Rowland, D. JESS, a Joint Expert Speciation System – VI: thermodynamically-consistent standard Gibbs energies of reaction for aqueous solutions. New J. Chem. 42, 7617–7629, DOI: 10.1039/C7NJ03597G (2018).

28. Anderson, G. M. Thermodynamics of Natural Systems (Cambridge University Press, Cambridge, 2005), 2nd edn.

29. Zhang, H. et al. Biosynthetic energy cost for amino acids decreases in cancer evolution. Nat. Commun. 9, 4124, DOI: 10.1038/s41467-018-06461-1 (2018).

30. Amend, J. P. & LaRowe, D. E. Mini-review: Demystifying microbial reaction energetics. Environ. Microbiol. 21, 3539–3547, DOI: 10.1111/1462-2920.14778 (2019).

31. Dick, J. M. & Shock, E. L. Calculation of the relative chemical stabilities of proteins as a function of temperature and redox chemistry in a hot spring. PLOS One 6, e22782, DOI: 10.1371/journal.pone.0022782 (2011).

32. Walsh, C. T., Tu, B. P. & Tang, Y. Eight kinetically stable but thermodynamically activated molecules that power cell metabolism. Chem. Rev. 118, 1460–1494, DOI: 10.1021/acs.chemrev.7b00510 (2018).

33. Morowitz, H. J. A theory of biochemical organization, metabolic pathways, and evolution. Complexity 4, 39–53, DOI: 10.1002/(SICI)1099-0526(199907/08)4:6<39::AID-CPLX8>3.0.CO;2-2 (1999).

34. DeBerardinis, R. J. & Cheng, T. Q’s next: The diverse functions of glutamine in metabolism, cell biology and cancer. Oncogene 29, 313–324, DOI: 10.1038/onc.2009.358 (2010).

35. Feist, A. M. et al. A genome-scale metabolic reconstruction for *Escherichia coli* K-12 MG1655 that accounts for 1260 ORFs and thermodynamic information. Mol. Syst. Biol. 3, 121, DOI: 10.1038/msb4100155 (2007).

36. Reeves, E. P., McDermott, J. M. & Seewald, J. S. The origin of methanethiol in midocean ridge hydrothermal fluids. Proc. Natl. Acad. Sci. 111, 5474–5479, DOI: 10.1073/pnas.1400643111 (2014).

37. Ooka, H., McGlynn, S. E. & Nakamura, R. Electrochemistry at deep-sea hydrothermal vents: Utilization of the thermodynamic driving force towards the autotrophic origin of life. ChemElectroChem 6, 1316–1323, DOI: 10.1002/celc.201801432 (2019).

38. Lindsay, M. R. et al. Subsurface processes influence oxidant availability and chemoautotrophic hydrogen metabolism in Yellowstone hot springs. Geobiology 16, 674–692, DOI: 10.1111/gbi.12308 (2018).

39. Ley, R. E. et al. Unexpected diversity and complexity of the Guerrero Negro hypersaline microbial mat. Appl. Environ. Microbiol. 72, 3685–3695, DOI: 10.1128/AEM.72.5.3685-3695.2006 (2006).

40. Swingley, W. D. et al. Coordinating environmental genomics and geochemistry reveals metabolic transitions in a hot spring ecosystem. PLOS One 7, e38108, DOI: 10.1371/journal.pone.0038108 (2012).

41. Boyer, G. M., Schubotz, F., Summons, R. E., Woods, J. & Shock, E. L. Carbon oxidation state in microbial polar lipids suggests adaptation to hot spring temperature and redox gradients. Front. Microbiol. 11, 229, DOI: 10.3389/fmicb.2020.00229 (2020).

42. Fortunato, C. S., Larson, B., Butterfield, D. A. & Huber, J. A. Spatially distinct, temporally stable microbial populations mediate biogeochemical cycling at and below the seafloor in hydrothermal vent fluids. Environ. Microbiol. 20, 769–784, DOI: 10.1111/1462-2920.14011 (2018).

43. Havig, J. R., Raymond, J., Meyer-Dombard, D. R., Zolotova, N. & Shock, E. L. Merging isotopes and community genomics in a siliceous sinter-depositing hot spring. J. Geophys. Res. 116, G01005, DOI: 10.1029/2010JG001415 (2011).

44. Reveillaud, J. et al. Subseafloor microbial communities in hydrogen-rich vent fluids from hydrothermal systems along the Mid-Cayman Rise. Environ. Microbiol. 18, 1970–1987, DOI: 10.1111/1462-2920.13173 (2016).

45. Dupont, C. L. et al. Functional tradeoffs underpin salinity-driven divergence in microbial community composition. PLOS One 9, 1–9, DOI: 10.1371/journal.pone.0089549 (2014).

46. Youens-Clark, K. et al. iMicrobe: Tools and data-driven discovery platform for the microbiome sciences. GigaScience 8, giz083, DOI: 10.1093/gigascience/giz083 (2019).

47. Asplund-Samuelsson, J. et al. Diversity and expression of bacterial metacaspases in an aquatic ecosystem. Front. Microbiol. 7, 1043, DOI: 10.3389/fmicb.2016.01043 (2016).

48. Satinsky, B. M. et al. Metagenomic and metatranscriptomic inventories of the lower Amazon River, May 2011. Microbiome 3, 39, DOI: 10.1186/s40168-015-0099-0 (2015).

49. Satinsky, B. M. et al. The Amazon continuum dataset: quantitative metagenomic and metatranscriptomic inventories of the Amazon River plume, June 2010. Microbiome 2, 17, DOI: 10.1186/2049-2618-2-17 (2014).

50. Eiler, A. et al. Productivity and salinity structuring of the microplankton revealed by comparative freshwater metagenomics. Environ. Microbiol. 16, 2682–2698, DOI: 10.1111/1462-2920.12301 (2014).

51. Vavourakis, C. D. et al. Metagenomic insights into the uncultured diversity and physiology of microbes in four hypersaline soda lake brines. Front. Microbiol. 7, 211, DOI: 10.3389/fmicb.2016.00211 (2016).

52. Ghai, R. et al. New abundant microbial groups in aquatic hypersaline environments. Sci. Reports 1, 135, DOI: 10.1038/srep00135 (2011).

53. Fernandez, A. B. et al. Metagenome sequencing of prokaryotic microbiota from two hypersaline ponds of a marine saltern in Santa Pola, Spain. Genome Announc. 1, 6, DOI: 10.1128/genomea.00933-13 (2013).

54. Kimbrel, J. A. et al. Microbial community structure and functional potential along a hypersaline gradient. Front. Microbiol. 9, 1492, DOI: 10.3389/fmicb.2018.01492 (2018).

55. Dick, J. M. Average oxidation state of carbon in proteins. J. Royal Soc. Interface 11, 20131095, DOI: 10.1098/rsif.2013.1095 (2014).

56. Rhodes, M. E., Fitz-Gibbon, S. T., Oren, A. & House, C. H. Amino acid signatures of salinity on an environmental scale with a focus on the Dead Sea. Environ. Microbiol. 12, 2613–2623, DOI: 10.1111/j.1462-2920.2010.02232.x (2010).

57. Leuko, S., Raftery, M. J., Burns, B. P., Walter, M. R. & Neilan, B. A. Global protein-level responses of *Halobacterium salinarum* NRC-1 to prolonged changes in external sodium chloride concentrations. J. Proteome Res. 8, 2218–2225, DOI: 10.1021/pr800663c (2009).

58. Fränzel, B. et al. Adaptation of *Corynebacterium glutamicum* to salt-stress conditions. Proteomics 10, 445–457, DOI: 10.1002/pmic.200900482 (2010).

59. Hahne, H. et al. A comprehensive proteomics and transcriptomics analysis of *Bacillus subtilis* salt stress adaptation. J. Bacteriol. 192, 870–882, DOI: 10.1128/JB.01106-09 (2010).

60. Zhang, Y. et al. Quantitative proteomics reveals membrane protein-mediated hypersaline sensitivity and adaptation in halophilic *Nocardiopsis xinjiangensis*. J. Proteome Res. 15, 68–85, DOI: 10.1021/acs.jproteome.5b00526 (2016).

61. Lin, J., Liang, H., Yan, J. & Luo, L. The molecular mechanism and post-transcriptional regulation characteristic of *Tetragenococcus halophilus* acclimation to osmotic stress revealed by quantitative proteomics. J. Proteomics 168, 1–14, DOI: 10.1016/j.jprot.2017.08.014 (2017).

62. Lee, T. et al. Salt stress affects global protein expression profiles of extracellular membrane-derived vesicles of *Listeria monocytogenes*. Microb. Pathog. 115, 272–279, DOI: 10.1016/j.micpath.2017.12.071 (2018).

63. Jevtić, v. et al. The response of *Haloferax volcanii* to salt and temperature stress: A proteome study by label-free mass spectrometry. Proteomics 19, 1800491, DOI: 10.1002/pmic.201800491 (2019).

64. Bolker, B. M. Ecological Models and Data in R (Princeton University Press, Princeton, NJ, 2008).

65. Venables, W. N. & Ripley, B. D. Modern Applied Statistics with S (Springer, New York, 2002), 4th edn. http://www.stats.ox.ac.uk/pub/MASS4.

66. The UniProt Consortium. UniProt: A worldwide hub of protein knowledge. Nucleic Acids Res. 47, D506–D515, DOI: 10.1093/nar/gky1049 (2019).

67. de Goffau, M. C., van Dijl, J. M. & Harmsen, H. J. M. Microbial growth on the edge of desiccation. Environ. Microbiol. 13, 2328–2335, DOI: 10.1111/j.1462-2920.2011.02496.x (2011).

68. Simon, H. M., Smith, M. W. & Herfort, L. Metagenomic insights into particles and their associated microbiota in a coastal margin ecosystem. Front. Microbiol. 5, 466, DOI: 10.3389/fmicb.2014.00466 (2014).

69. Trigos, A. S., Pearson, R. B., Papenfuss, A. T. & Goode, D. L. Altered interactions between unicellular and multicellular genes drive hallmarks of transformation in a diverse range of solid tumors. Proc. Natl. Acad. Sci. 114, 6406–6411, DOI: 10.1073/pnas.1617743114 (2017).

70. Keegan, K. P., Glass, E. M. & Meyer, F. MG-RAST, a metagenomics service for analysis of microbial community structure and function. In Martin, F. & Uroz, S. (eds.) Microbial Environmental Genomics (MEG), 207–233, DOI: 10.1007/978-1-4939-3369-3_13 (Springer, New York, 2016).

71. Kopylova, E., Noé, L. & Touzet, H. SortMeRNA: Fast and accurate filtering of ribosomal RNAs in metatranscriptomic data. Bioinformatics 28, 3211–3217, DOI: 10.1093/bioinformatics/bts611 (2012).

72. Rho, M., Tang, H. & Ye, Y. FragGeneScan: Predicting genes in short and error-prone reads. Nucleic Acids Res. 38, e191, DOI: 10.1093/nar/gkq747 (2010).

73. R Core Team. R: A Language and Environment for Statistical Computing. R Foundation for Statistical Computing, Vienna, Austria (2019).

74. O’Leary, N. A. et al. Reference sequence (RefSeq) database at NCBI: current status, taxonomic expansion, and functional annotation. Nucleic Acids Res. 44, D733–D745, DOI: 10.1093/nar/gkv1189 (2016).

75. Kyte, J. & Doolittle, R. F. A simple method for displaying the hydropathic character of a protein. J. Mol. Biol. 157, 105–132, DOI: 10.1016/0022-2836(82)90515-0 (1982).

76. Bjellqvist, B. et al. The focusing positions of polypeptides in immobilized pH gradients can be predicted from their amino acid sequences. Electrophoresis 14, 1023–1031, DOI: 10.1002/elps.11501401163 (1993).

77. Bjellqvist, B., Basse, B., Olsen, E. & Celis, J. E. Reference points for comparisons of twodimensional maps of proteins from different human cell types defined in a pH scale where isoelectric points correlate with polypeptide compositions. Electrophoresis 15, 529–539, DOI: 10.1002/elps.1150150171 (1994).

78. Gasteiger, E. et al. Protein identification and analysis tools on the ExPASy server. In Walker, J. M. (ed.) The Proteomics Protocols Handbook, 571–607, DOI: 10.1385/1-59259-890-0:571 (Humana Press, Totowa, NJ, 2005).

